# Dual functions of Intraflagellar Transport Protein IFT20 in spermiogenesis: formation of sperm flagella and removal of cytoplasm by autophagy

**DOI:** 10.1101/051219

**Authors:** Zhengang Zhang, Wei Li, Yong Zhang, Ling Zhang, Maria E Teves, Hong Liu, Junpin Liu, Jerome F Strauss, Gregory J Pazour, James A Foster, Rex A. Hess, Zhibing Zhang

**Affiliations:** Department of Gastroenterology, Tongji Hospital, Tongji Medical College, Huazhong University of Science and Technology, Wuhan, Hubei, China, 430030; Department of Obstetrics and Gynecology, Virginia Commonwealth University, Richmond, VA, 23298; Department of Dermatology, Tongji Hospital, Tongji Medical College, Huazhong University of Science and Technology, Wuhan, Hubei, China, 430030; School of Public Health, Wuhan University of Science and Technology, Wuhan, Hubei, 430065; Wuhan Hospital for the Prevention and Treatment of Occupational Diseases; Program in Molecular Medicine, University of Massachusetts Medical School, Worcester, MA 01605; Department of Biology, Randolph-Macon College, Ashland, VA 23005; Comparative Biosciences, College of Veterinary Medicine, University of Illinois, 2001 S. Lincoln, Urbana, IL 61802-6199.

**Keywords:** Intraflagellar transport, spermatogenesis, flagellogenesis, retained cytoplasm, autophagy

## Abstract

Intraflagellar transport (IFT) is a conserved mechanism thought to be essential for the assembly and maintenance of cilia and flagella. However, little is known about mammalian sperm flagella formation. To fill this gap, we disrupted the *Ift20* gene in male germ cells. Homozygous mutant mice were infertile with significantly reduced sperm counts and motility. In addition, abnormally shaped elongating spermatid heads and bulbous round spermatids were found in the lumen of the seminiferous tubules. Electron microscopy revealed increased cytoplasmic vesicles, fiber-like structures, abnormal accumulation of mitochondria and decreased mature lysosomes. The few developed sperm had disrupted axoneme and retained cytoplasmic lobe components on the flagella. ODF2 and SPAG16L, two sperm flagella proteins failed to be incorporated into sperm tails of the mutant mice. Expression levels of an autophagy core protein that associates with IFT20, Atg16, were significantly reduced in the testis of the *Ift20* mutant mice. Our studies suggest that IFT20 is essential for spermiogenesis in mice, and it plays a role in sperm flagella formation, and removing excess cytoplasmic components by regulating autophagy core proteins.

## Summary Statement.

We explored the role of IFT20 in male germ cell development, and discovered that IFT20 plays a role in building the sperm flagella and the disposal of cytoplasmic components by autophagy.

## Introduction

Cilia are cell surface organelles that are ubiquitously present on almost all vertebrate cells, at some point during development and differentiation (Brown and Witman, 2014). They are formed by a process that requires intraflagellar transport (IFT), a bidirectional transport system directed by IFT protein complexes and motors (Pedersen and Rosenbaum, 2008). IFT involves movement of large protein complexes called IFT particles from the cell body to the ciliary tip (anterograde transport), followed by their return to the cell body (retrograde transport) (Blacque et al., 2008). It was first described in *Chlamydomonas* (Kozminski et al., 1993). IFT protein complexes have been named complex A and complex B, containing about 20 IFT proteins in *Chlamydomonas* (Fort and Bastin, 2014). Complex A is thought to be composed of six subunits, and complex B contains 16 known proteins including IFT20. IFT proteins are typically found along the ciliary shaft and at the base (Mizuno et al., 2012), and the IFT complexes serve as adaptors to mediate contacts between cargo proteins and motors (Follit et al., 2009). Mutations in both complex A and B IFT genes result in ciliogenesis defects and human diseases (Murcia et al., 2000; Huangfu and Anderson, 2005; Badano et al., 2006; Tran et al., 2008; Cortellino et al., 2009; Stottmann et al., 2009; Friedland-Little et al., 2011; Hildebrandt et al., 2011; Huber and Cormier-Daire, 2012).

IFT20 is the smallest IFT protein. The role of mouse IFT20 has been studied *in vitro* and *in vivo*. Among all identified mouse IFT proteins, IFT20 is the only one present in the Golgi body, and the localization is dependent on GMAP210. In living cells, fluorescently tagged IFT20 is highly dynamic and moves between the Golgi complex and the cilium as well as along ciliary microtubules; it functions in the delivery of ciliary membrane proteins from the Golgi complex to the cilium (Follit et al., 2006; Follit et al., 2008).

Disruption of the gene globally by a conventional knockout strategy resulted in embryonic lethality, demonstrating that IFT20 is required for development (Keady et al., 2011). The role of IFT20 in individual cells/tissues has been explored using floxed *Ift20* mice crossed to several tissue specific Cre transgenic mice. A variety of phenotypes relating to cilia dysfunction have been identified (Jonassen et al., 2008; May-Simera et al., 2015). IFT20 was also found to be expressed in non-ciliated T cells and to have functions in immune synapse formation (Finetti et al., 2009; Finetti et al., 2014; Yuan et al., 2014). Recent studies demonstrated that part of the molecular machinery involved in ciliogenesis also participates in autophagy (Pampliega et al., 2013). Autophagy is the catabolic process that involves degradation of unnecessary cellular components through the actions of lysosomes (Kobayashi, 2015). This process is mediated by the autophagy-related genes (Atg) and their associated enzymes. IFT20 associates with autophagy core protein Atg16L, and directs normal autophagy to clear redundant cytoplasmic components (Pampliega et al., 2013).

IFT proteins usually associate with other proteins to form a complex in order to conduct their functions. Besides Atg16L and GMAP210, IFT20 was reported to associate with several other proteins, including IFT57, a kinesin II subunit, KIF3B, and CCDC41 (Baker et al., 2003; Joo et al., 2013).

Even though IFT20 has been shown to play roles in both ciliated and non-ciliated cells, it is not clear if it also controls sperm tail formation. Of all the mammalian cells, sperm have the longest flagella. IFT20 is expressed in male gem cells. It is present in the Golgi complex, manchette, and the basal bodies of differentiating germ cells. These are key structures in ciliogenesis (Sironen et al., 2010). In male germ cells, it co-localizes with sperm flagella 2 (SPEF2), a key spermatogenesis regulator (Sironen et al., 2010; Sironen et al., 2011; Sironen et al., 2012). However, the exact role of IFT20 in sperm development remains unknown. In order to elucidate the function of IFT20 in the testis, particularly in male germ cell development, we generated a male germ cell specific *Ift20* knockout mouse model by crossing the floxed *Ift20* mice to *Stra8-iCre* transgenic mice. Our studies demonstrated that IFT20 is essential for spermiogenesis and male fertility. As expected, sperm flagella formation was disrupted in the conditional *Ift20* mutant mice. The unexpected finding was that some sperm that completed all phases of spermatogenesis retained the cytoplasmic lobe residue, which was associated with reduced expression of the autophagy core protein Atg16 in the testis. These findings suggest dual functions of IFT20 in male germ cell development: directing sperm flagella formation by IFT mechanisms and helping to remove residual cytoplasmic components by autophagy.

## Results

### Mouse IFT20 protein is highly abundant in the testis and its expression is up-regulated during spermiogenesis

By using Western Blot analysis, expression levels of IFT20 protein in various tissues were analyzed, including brain, lung, kidney, spleen and testis. IFT20 protein was only detected in the testis using the Pico ECL system (Fig. 1A). Next, we examined the testicular expression levels during germ cell development. The expression of IFT20 protein was first visible at post-natal day 16, and increased significantly at days 30 and 42, when germ cells enter the condensation/elongation stage in the first wave of spermatogenesis (Fig. 1B).

**Figure 1.**
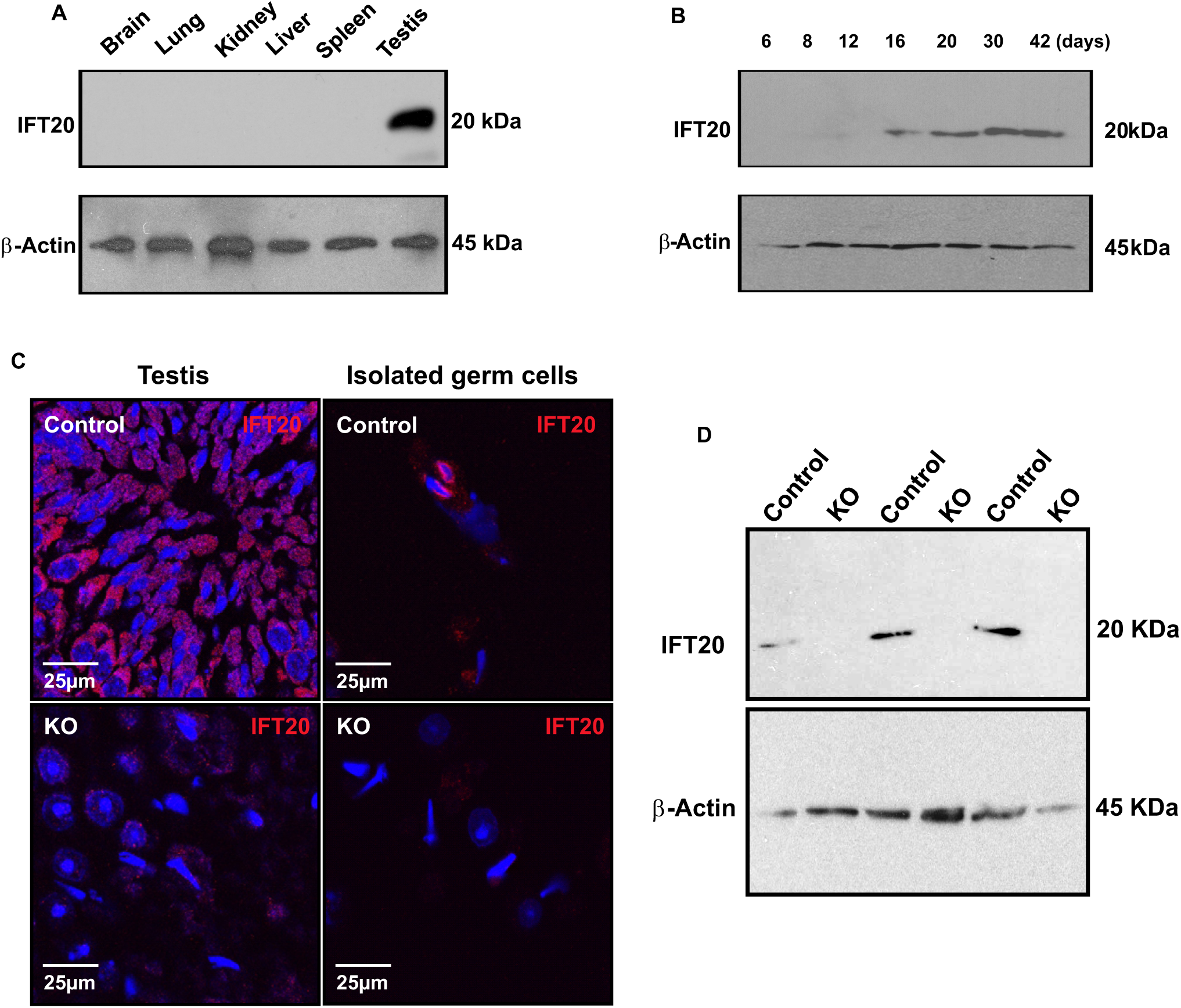
IFT20 protein expression in the wild type and conditional mutant mice. A. Tissue distribution of mouse IFT20 tissue expression. Notice that IFT20 is highly expressed in the testis. B: IFT20 expression levels during the first wave of spermatogenesis. Notice that its expression is significantly increased at day 16 of birth, and further increased at day 30 of birth when germ cells enter spermiogenesis. C. Examination of IFT20 protein expression by immunofluorescence staining on testicular sections (left) and isolated germ cells (Right). Notice that strong IFT20 signals (red) were observed in the control samples (upper). However, almost no specific IFT20 signals were observed in the knockout samples (lower). D. Analysis of testicular IFT20 expression by Western blot using the Pico system. Notice that IFT20 was expressed in the three control mice. However, no specific signal was observed in the three mutant mice analyzed.

### Generation of male germ cell specific *Ift20* conditional knockout mice

To study the role of IFT20 in mouse spermatogenesis, particularly spermiogenesis when germ cells form flagella, we generated male germ cell specific conditional knockout mice by crossing the *Stra8-iCre* males with *Ift20*^*flox/flox*^ females, following the breeding strategy shown in Fig. S1A. The resulting mice were genotyped by PCR (Fig. S1B) using the specific primers as indicated in Materials and Methods and previous publications (Jonassen et al., 2008). *Stra8-iCre* expression started in a proportion of spermatogonia around P3 or P4 and continued to increase with male germ cell development from spermatogonia to spermatocytes and then spermatids. The iCre-mediated excision reached full efficiency only in pachytene spermatocytes and spermatids in adult testes, whereas levels of iCre decreased to the minimum in spermatogonia in the adult testes (Bao et al., 2013). To examine IFT20 protein expression in the mutant mice, immunofluorescence staining and Western blotting were conducted. The control mice showed robust IFT20 signals in both testicular sections (Fig. 1C, left, upper panel) and isolated germ cells (Fig. 1C, right, upper panel). However, specific IFT20 signal was barely detectable in the testicular sections (Fig. 1C, left, lower panel) and isolated germ cells (Fig. 1C, right, lower panel). Western blot analysis confirmed that no specific IFT20 signal was detectable in the testes of the *Ift20* mutant mice when the less sensitive Pico system was used (Fig. 1D). However, when a highly sensitive Femto system was used, very low levels of IFT20 were still detected in the mutant mice (Fig. S2).

### Homozygous mutant adult *Ift20* males are grossly normal but infertile

All mutant mice survived to adulthood and did not show any gross abnormalities. To test fertility, two to three month-old *Ift20* mutant males and controls were bred with three to four month-old WT females. All 12 controls exhibited normal fertility, and sired normal sized litters after one month of breeding. However, none of the 14 *Ift20* mutant males tested produced any litters after three months of breeding (Table 1), even though these mice showed normal sexual behavior, and vaginal plugs were observed in the females that were bred with the *Ift20* mutant males.

**Table 1.**
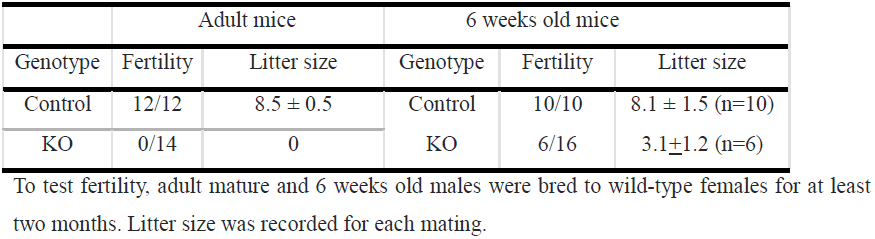
Fertility, fecundity of control and conditional *Ift20* mutant mice.

| Adult mice |  |  | 6 weeks old mice |  |  |
| --- | --- | --- | --- | --- | --- |
| Genotype | Fertility | Litter size | Genotype | Fertility | Litter size |
| Control | 12/12 | $8.5 \pm 0.5$ | Control | 10/10 | $8.1 \pm 1.5$ (n=10) |
| KO | 0/14 | 0 | KO | 6/16 | $3.1 \pm 1.2$ (n=6) |

To test fertility, adult mature and 6 weeks old males were bred to wild-type females for at least two months. Litter size was recorded for each mating.

A fertility test was also performed with a group of control and homozygous conditional *Ift20* mutant mice at six weeks of age to examine the mutant model during the first wave of spermatogenesis. Interestingly, even though all adult mutant males were infertile, 37.5% of the 6 weeks old conditional *Ift20* mutant males (six of sixteen) sired pups. However, the litter sizes (2-5 pups per litter) were significantly smaller than the controls at the same age (7-11 pups per litter). The six homozygous mutant males never sired pups again after the first litter (Table 1).

### Abnormal sperm morphology, significantly reduced sperm count and sperm motility

To determine sperm factors in the infertility phenotype, sperm morphology, number, and motility were examined in control and *Ift20* mutant mice. Sperm were collected from cauda epididymides of the control and conditional *Ift20* mutant mice. Under light microscopy, sperm density was remarkably lower in the mutant mice compared to that of the controls under the same dilution. The tails of many sperm were kinked or shortened (Fig. 2A). By SEM, sperm from the control mice had long and smooth tails with normally condensed heads (Fig. 2B-a). However, multiple abnormalities were observed in sperm of the mutant mice, including short and kinked tails that were also observed under light microscopy (Fig. 2Bb-d). In addition, some sperm showed round and swollen heads (Fig. 3Bc, e, f). Some sperm with kinked tails had retained cytoplasm in the kinked area (Fig. 2Bd). Sperm numbers in the mutant mice were significantly lower than in controls (Fig. 2C). Movies were taken from swim-out sperm. Most control sperm displayed vigorous flagellar activity and progressive long track forward movement (Movie. S1 and Fig. 2D). However, only a small percentage of the mutant sperm were motile and showed forward motility, and flagellar movement was limited to a slow, erratic waveform with a very low amplitude (Movie. S2 and Fig. 2D). Even if some sperm were motile, their motility would have been significantly less than the controls (Fig. 2E).

**Figure 2.**
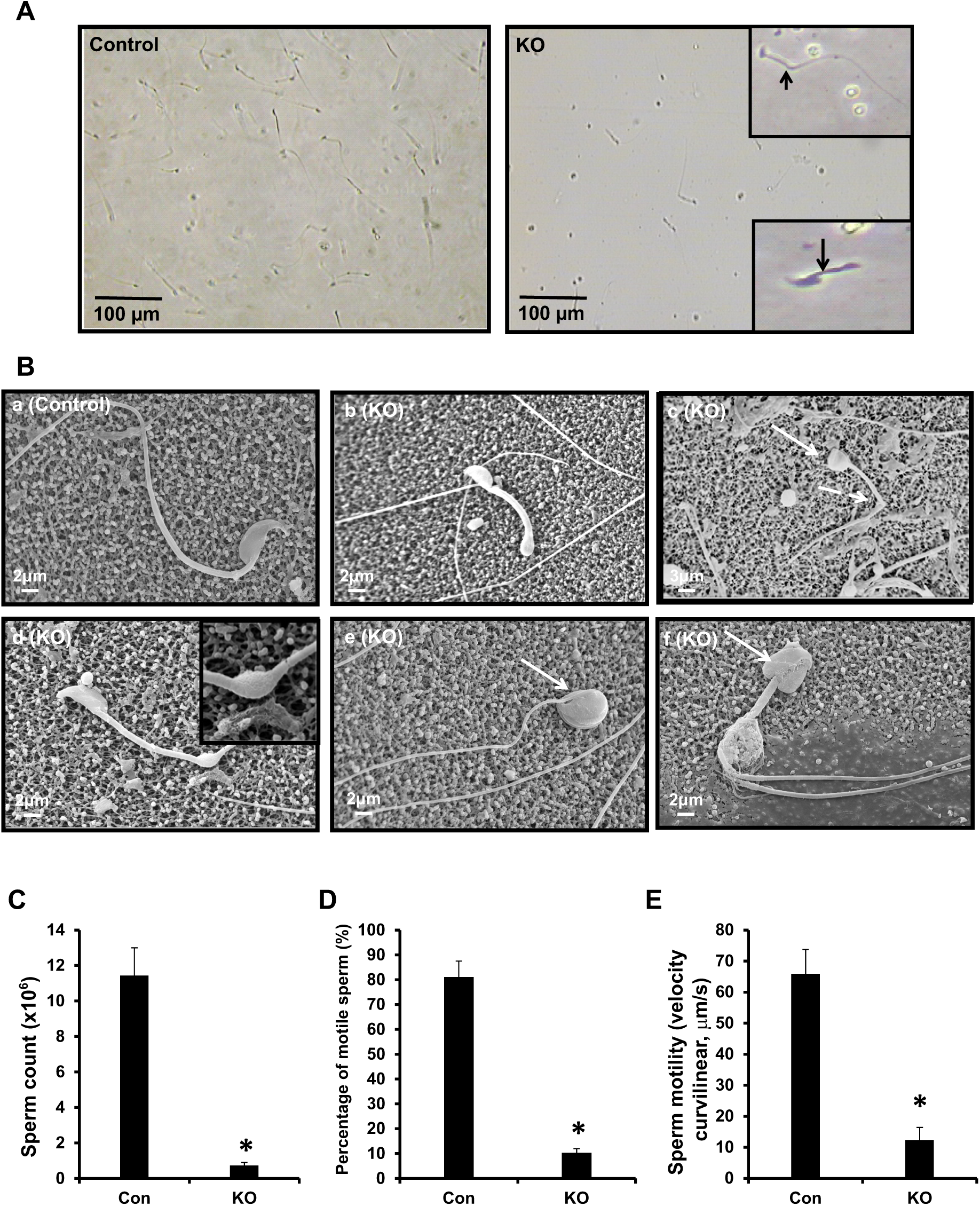
Disruption of *Ift20* gene in germ cells leads to abnormal sperm morphology associated with significant reduced sperm count and sperm motility. A. Examination of epididymal sperm by light microscopy. Under the same dilution, compared to the control (left panels), very few sperm were seen in the conditional *Ift20* mutant mice (right panels). Sperm from the mutant mice often had shorter (arrow) or kinked (arrow head) tails (inserts in the right panels). B. Examination of epididymal sperm by SEM. a: Representative image of epididymis sperm with normal morphology from a control mouse shows the sperm with a long and smooth tail and condensed head. b to f: Representative images of epididymal sperm from a mutant mouse. A variety of abnormal sperm morphologies are observed, with short (b) and kinked tails (c, d); some with round heads (c, e, f). The insert in “d” shows excess retention of cytoplasm. The white arrows point to sperm with round heads, the dashed arrows point to bend tails. C. The sperm count was significantly reduced in the mutant mice; D. The percentage of motile sperm was significantly lower in the mutant mice; E. Sperm VCL was significantly reduced in the mutant mice compared to the controls (*p<0.05).

**Figure 3.**
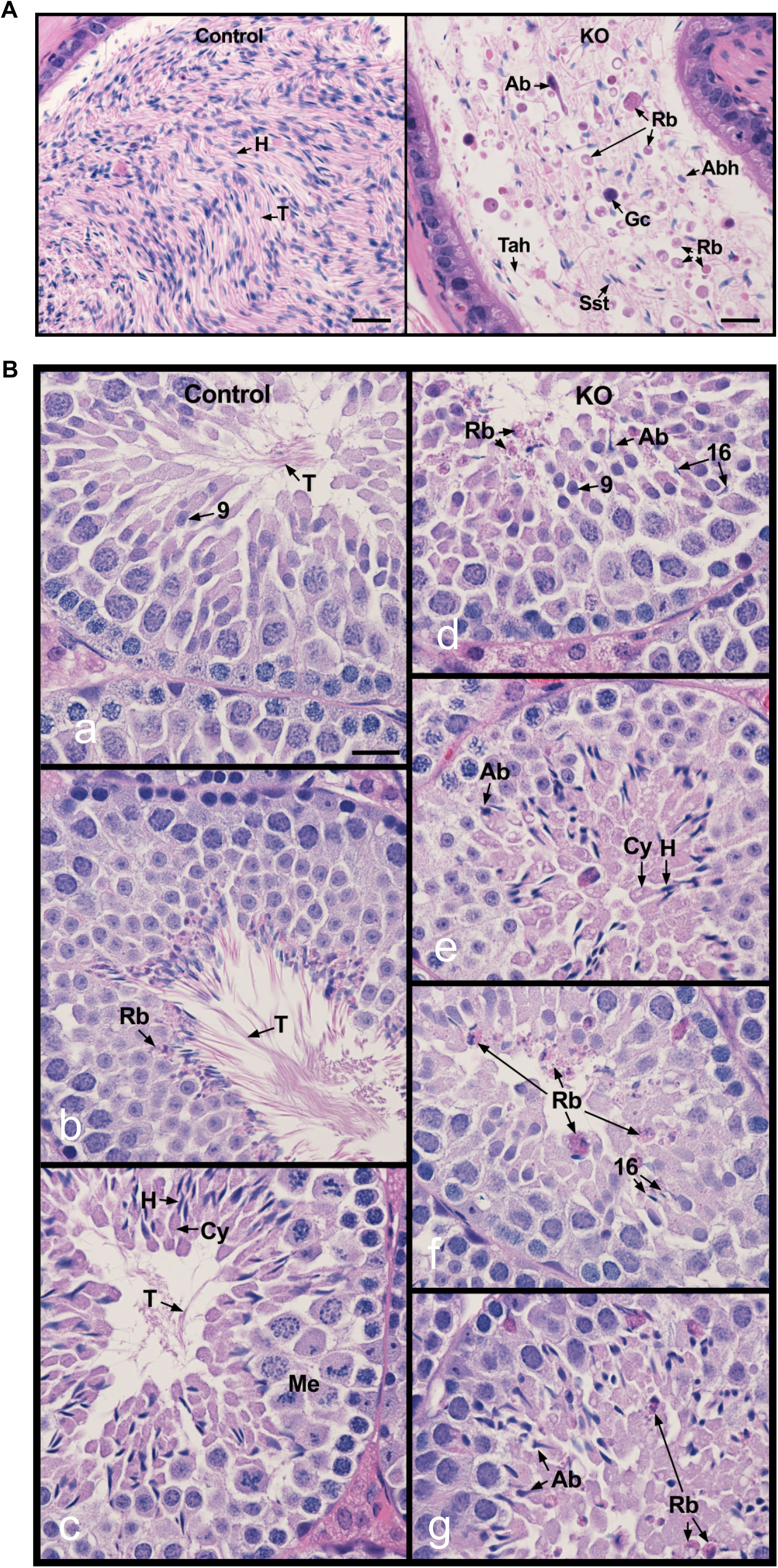
Examination of histology of cauda epididymis (A) and testis (B) of control and conditional *Ift20* mutant mice. A. Control showing highly concentrated epididymal sperm with aligned sperm tails (T) and normal sperm heads (H) (left panel). B. IFT20 knockout (right) showing a very low concentration of sperm, with high incidence of sperm abnormalities (Ab), sloughed round germ cells (Gc), residual bodies (Rb), sperm with short tails (Sst) and sperm tails without heads (Tah). Bars = 20 mm. B. Testes of control (a-c) and IFT20 knockout mice (IFT20 KO; d-g). a. Stage IX showing normal step 9 spermatids with long tails (T) extending into the lumen. b. Stage VIII showing mature sperm with long tails (T) being released into the lumen. Residual bodies (Rb) of left over cytoplasm are being formed and phagocytized by Sertoli cells. c. Stage XII showing spermatocytes in meiotic division (Me), step 12 elongating spermatids with head (H) of condensed chromatin and bulges of cytoplasm (Cy). The spermatid tails (T) are long and extend into the lumen. d. Stage IX showing abnormal step 16 elongated spermatids (Ab) that were not released into the lumen and residual bodies (Rb) that are being sloughed into the lumen. e. Stage I-III showing abnormal elongated spermatids (Ab). The elongated spermatids show some normal heads (H) and cytoplasm (Cy), but their tails do not appear in the lumen (*). f. Stage X showing failure of spermiation, with step 16 spermatid heads remaining in the seminiferous epithelium. Residual bodies (Rb) remain at the lumen, rather than being phagocytized. g. Stage XI showing abnormal elongating spermatids (Ab) and sloughed residual bodies (Rb) remaining from prior stages. Bar = 20 mm for all photos.

### Very few well-developed epididymal sperm and abnormal spermiogenesis in the conditional *Ift20* mutant mice

Histology of cauda epididymides of adult control and conditional *Ift20* mutant mice was examined. The control mice showed highly concentrated epididymal sperm with aligned sperm tails and normal sperm heads (Fig. 3A, left). However, *Ift20* mutant mice showed a very low concentration of sperm, with high incidence of sperm abnormalities, sloughed round germ cells, residual bodies, and sperm with short tails or sperm tails without heads (Fig. 3A, right).

To understand the testicular factors that contribute to low sperm counts and abnormal sperm morphology, testes of the control (Fig. 3Ba-c) and *Ift20* mutant mice (Fig. 3Bd-g) were examined. There was no difference between testis weight/body weight between controls and mutant mice (Fig. S3). Testis histology was further investigated (Fig. 3B and Fig. S4). The control mice showed normal completion of spermatogenesis. Stage IX showed normal step 9 spermatids with long tails (T) extending into the lumen (Fig. 3Ba). Stage VIII showed mature sperm with long tails (T) being released into the lumen. Residual bodies (Rb) of leftover cytoplasm were being formed and phagocytized by Sertoli cells (Fig. 3Bb). Stage XII showed spermatocytes in meiotic division (Me), step 12 elongating spermatids with heads (H) of condensed chromatin and bulges of cytoplasm (Cy). The spermatid tails (T) were long and extend into the lumen (Fig. 3Bc). In the mutant testis, Stage IX showed abnormal step 16 elongated spermatids (Ab) that were not released into the lumen and residual bodies (Rb) that were being sloughed into the lumen (Fig. 3Bd). In mutant Stages I-III (Fig. 3Be), abnormal elongated spermatids (Ab) were observed along with some normal heads (H) and cytoplasm (Cy), but spermatid tails did not appear in the lumen (*). Mutant Stage X (Fig. 3Bf) showed failure of spermiation, with step 16 spermatid heads remaining in the seminiferous epithelium and residual bodies (Rb) remain at the lumen, rather than being phagocytized. Mutant Stage XI showed abnormal elongating spermatids (Ab) and sloughed residual bodies (Rb) remaining from prior stages (Fig. 3Bg).

### Ultrastructural changes in the sperm and testis of the conditional *Ift*20 mutant mice

TEM was conducted to examine ultrastructure changes in epididymal sperm (Fig. 4A, 4B, Fig. S5, 6) and testis (Fig. 4C, Fig. S7) of control and the *Ift*20 mutant mice. Sperm cells were the predominant cell type in the control mouse seminiferous tubule lumen (Fig. 4A, left panel). However, few sperm were present in the mutant mice, and significant numbers of residual bodies were present (Fig. 4A, right panel, and Fig. S5). Control mice showed normal sperm structure (Figure 4B, let upper panel). However, abnormal axonemal structures were frequently discovered in the mutant mice, including a disrupted “9+2” pattern (Fig. 4B, right upper panel), missing ODFs in the axoneme (Fig. 4B, lower, left panel). Some sperm retained excess retention of cytoplasm (Fig. 4B, lower, right panel, and Fig. S6). High magnification images of the residual bodies demonstrated that they contained mitochondria, fibrous sheath materials, microtubules, outer dense fibers, and membranes that normally would have been part of the assembled flagella (Fig. 4C).

**Figure 4.**
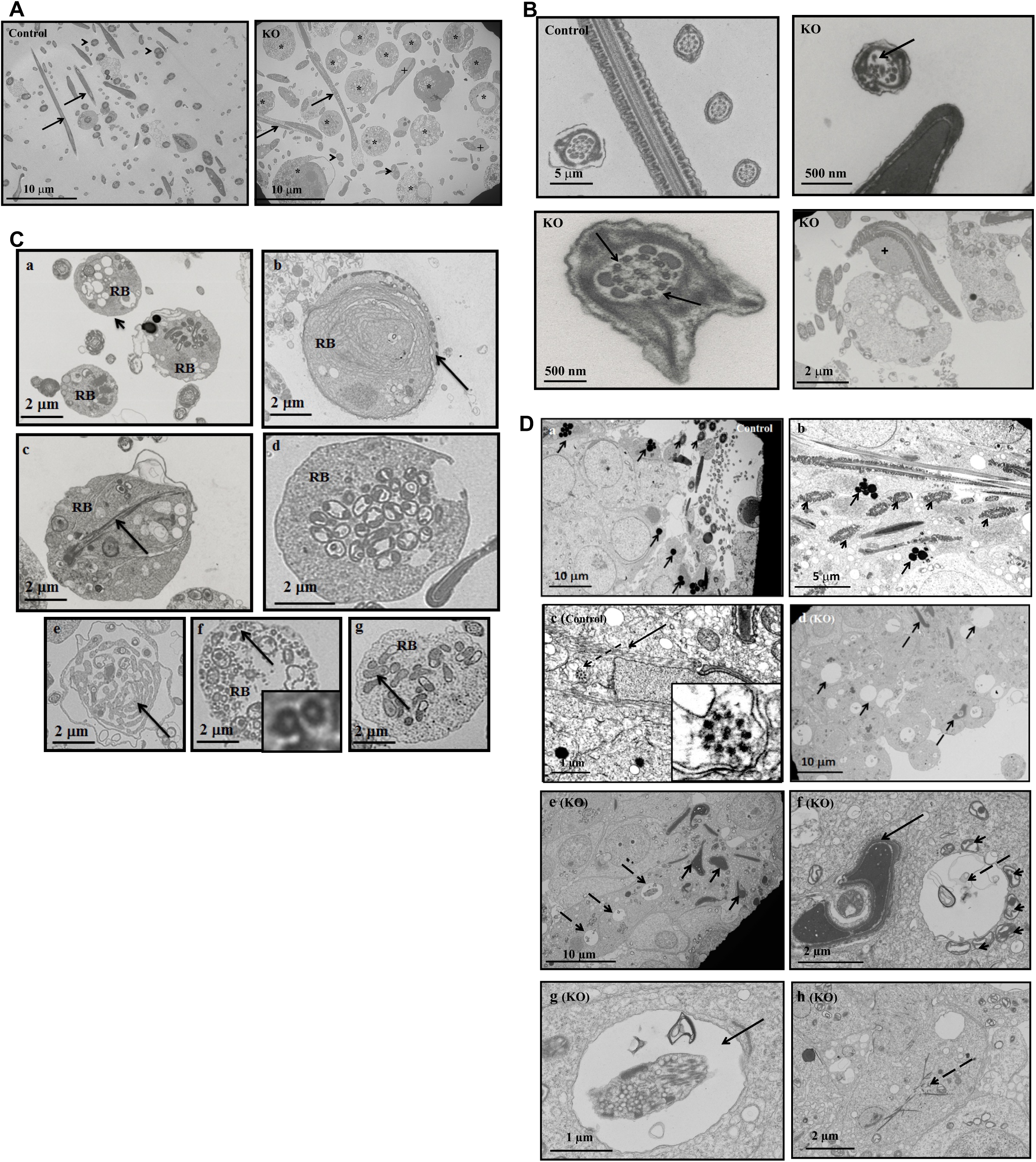
Ultrastructure of sperm from epididymis and testis of control and conditional *Ift20* mutant mice. A. Low magnification images of cells collected from cauda epididymides of a control mouse and a conditional *Ift20* mutant mouse. Notice that sperm cells are the predominant cell type in the control mouse. However, few sperm were present in the mutant mouse and some of these sperm had excess cytoplasm (+). Also, significant numbers of residual bodies (*) were present. The arrows point to the sperm axoneme in longitudinal section and the arrow heads point to the sperm axoneme in cross-section. B. Representative high magnification images of sperm axonemes from control and conditional *Ift20* mutant mice. Notice that the sperm axonemes from the control mouse show normal architecture. However, multiple defects were frequently discovered in sperm from the mutant mice. The arrow points to an axoneme with disrupted “9+2” structure; the dashed arrows to an axoneme missing three ODFs; the “+” sign indicates sperm with excess retention of cytoplasm. C. Representative high magnification images of residual bodies obtained from cauda epididymides of conditional *Ift20* mutant mice. Residual bodies are normally phagocytosed by Sertoli cells during spermiation and they contain components from sperm that failed to assemble correctly; some identifiable structures include mitochondria (a, d, g); fibrous sheath (c), microtubules (f); outer dense fibers (g); and membranes (b, e). D. Ultrastructure of the testis of control and conditional *Ift20* mutant mice. Panel “a” shows a representative image of the lumen area of a seminiferous tubule from a control mouse. Notice that numbers of sperm axoneme were present (arrow heads), and mature lysosomes (arrows) were easily seen. Panel “b” shows a higher magnification image of a seminiferous tubule from a control mouse. Numbers of developing axoneme (arrow heads) and mature lysosome (arrows) were observed. Panel “c” shows a developing spermatid. The arrow points to the manchette; the dashed arrow points to a developing axoneme. The zoomed-in image (the insert) shows the “9+2” structure of the developing axoneme. Panel “d” shows a representative image of the lumen area of a seminiferous tubule from a conditional *Ift20* mutant mouse. Axoneme structure can hardly be observed. Instead, a large number of large vacuoles (arrows) as well as deformed chromatin (dashed arrows) were discovered. Panel “e” shows a higher magnification image of a seminiferous tubule from a conditional *Ift20* mutant mouse. Neither axoneme nor mature lysosomes were present. Instead, degenerating chromatin (arrows) and vacuoles (dashed arrows) were throughout the seminiferous tubules. The arrow in “f” points to a deformed chromatin; the dashed arrow points to a large vacuole surrounded by many mitochondria (arrow heads). The arrows in “g” point to large vacuoles with various contents inside. The dashed arrow in “h” points to some fiber-like structures.

In the control testis, well-developed sperm were present in the lumen of seminiferous tubules, as indicated by the cross-section of normal axonemes. Dark lysosomes were seen in the cytoplasm of late step spermatids (Fig. 4D, a, b). The elongating spermatids from the control mice had normal developing manchette microtubules (Fig. 4D, c), as well as normally condensed chromatin (Fig. 4D). In the mutant testes, flagellar axonemes were rarely seen in seminiferous tubules and lumen (Fig. 4D, d). Large vacuoles and abnormally condensed chromatin were present in the tubule lumen and seminiferous tubules (Fig. 4D, e); some vacuoles were surrounded by mitochondria, some vacuoles had various contents inside; and some cells had fiber-like cytoplasmic structures (Figure 4D, f-h, and Fig. S7).

### Expression levels of an outer dense fiber component, ODF2 and a central apparatus component, SPAG16L are normal, but they are not assembled into sperm flagella of the *Ift*20 mutant mice

To examine the expression of components of sperm flagella of control and conditional *Ift20* mutant mice, Western blotting and immunofluorescence staining were conducted targeting two representative proteins: ODF2, an outer dense fiber component, and SPAG16L, a central apparatus component. Western blotting analysis revealed that there was no difference in the testicular expression levels of both proteins between the control and *Ift*20 mutant mice (Fig. 5A). Immunofluorescence staining demonstrated that both ODF2 and SPAG16L were present in the developing flagella of elongating spermatids isolated from the control mice (left panels of Fig. 5B and Fig. 5C). However, even though the two proteins were expressed in the elongating spermatids isolated from the *Ift20* mutant mice, the proteins were still in the cytoplasm (right panels of Fig. 5B and Fig. 5C).

**Figure 5.**
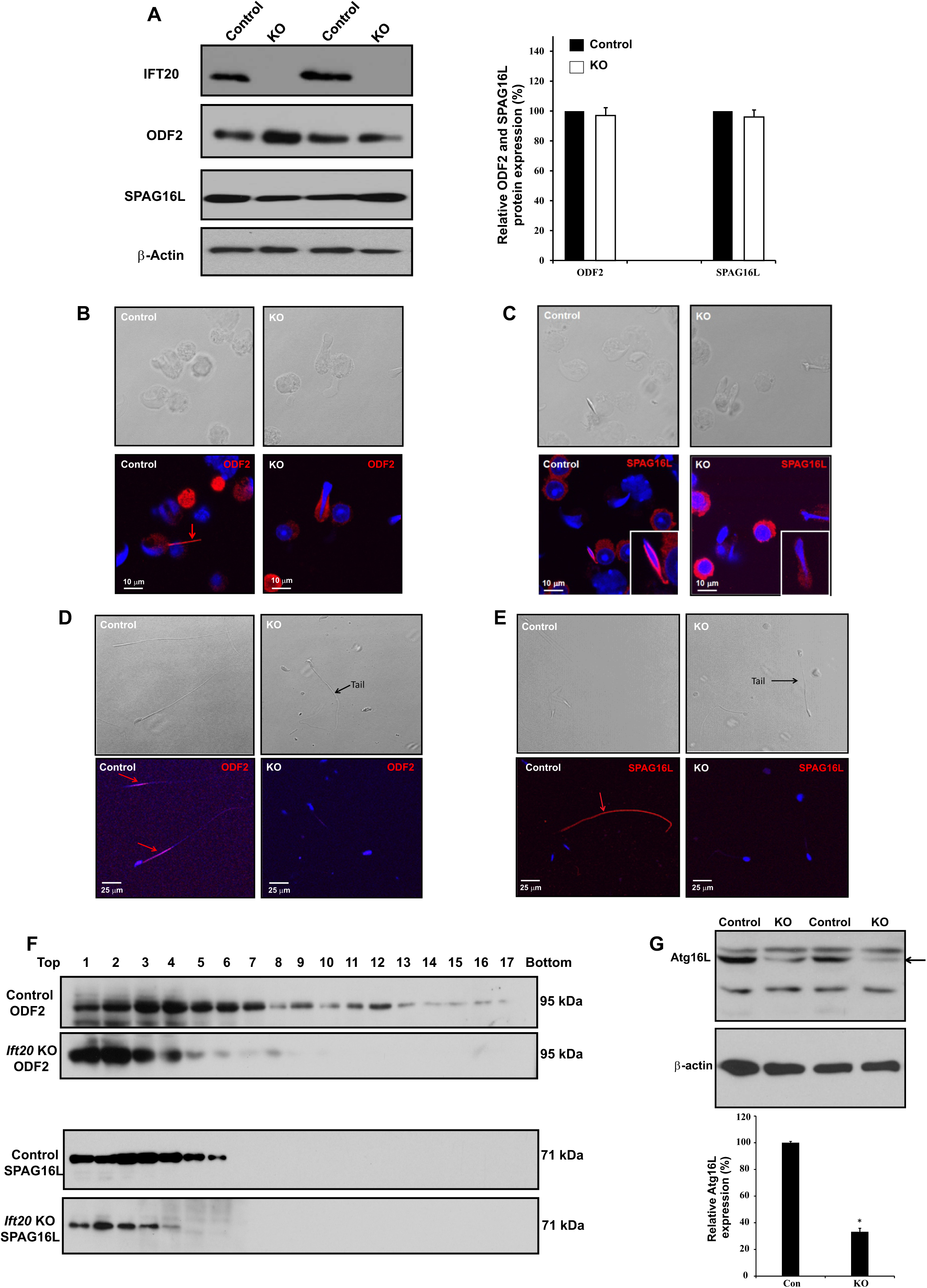
Expression and localization of sperm flagellar proteins and autophagy core protein Atg16L in the control and *Ift*20 mutant mice. A. Testicular expression levels of a major component of sperm tail outer dense fibers (ODF), ODF2, and a central apparatus protein, SPAG16L, are not changed in the *Ift*20 mutant mice. Left. Representative Western blot results using specific antibodies to examine testicular ODF2 and SPAG16L expression levels in control and *Ift*20 mutant mice. Right. Quantitation of relative ODF2 and SPAG16L expression from three control and three *Ift*20 mutant mice. There is no significant difference in expression levels between control and mutant mice. B. ODF2 and SPAG16L are not assembled into flagella correctly in the *Ift20* mutant mice. Representative image of localization of ODF2 in the elongating spermatids of a control mouse (left) and an *Ift20* mutant mouse (right). Specific signal (red) was detected in the developing tail of an elongating spermatid of the control mouse. However, ODF2 signal was still in the cytoplasm of an elongating spermatid of *Ift20* mutant mouse. C. Representative image of localization of SPAG16L in the elongating spermatids of a control mouse (left) and an *Ift20* mutant mouse (right). Specific signal (red) was detected in the developing tail of an elongating spermatid of the control mouse. However, SPAG16L was still in the cytoplasm of an elongating spermatid of *Ift20* mutant mouse. D. Representative image of localization of ODF2 in the epididymis sperm of a control mouse (left) and an *Ift20* mutant mouse (right). Specific signal (red) was observed in the principle piece of the control mouse. However, no specific signal was seen in *Ift20* mutant mouse. E. Representative image of localization of SPAG16L in the epididymis sperm of a control mouse (left) and an *Ift20* mutant mouse (right). Specific signal (red) was detected in whole sperm tail of the control mouse. However, SPAG16L signal was not observed in the few developed sperm of the *Ift20* mutant mouse. F. The distribution of testicular ODF2 and SPAG16L in sucrose gradient is altered in the *Ift20* mutant mice. Notice that both ODF2 and SPAG16L had a wider distribution in the gradients. However, distribution of the two proteins appeared to shift to the lighter fractions. G. Expression of autophagy core protein Atg16L is dramatically reduced in the testis of the conditional *Ift20* mutant mice. Left: Representative Western blot result of two control and two *Ift20* mutant mice. Notice that compared to the two control mice, testicular expression of ATG16L is lower in the two *Ift20* mutant mice. Right: Statistical analysis of relative expression of ATG16L normalized by β-actin from three control and three *Ift20* mutant mice. Relative ATG16L expression in the *Ift20* mutant mice is significantly lower than that in the control mice. * p<0.05.

Localization of the two proteins was also examined in epididymal sperm. In control mice, ODF2 was present in the flagellar midpiece (Fig. 5D, left panel) and SPAG16L was present along the entire flagellum (Fig. 5E, left panel). However, when a tail was formed in the *Ift20* mutant sperm, no ODF2 or SPAG16L protein was detected (right panels of Fig. 5D and Fig. 5E).

### Different distribution of ODF2 and SPAG16L in sucrose gradient between control and *Ift20* mutant mice

Distribution of testicular ODF2 and SPAG16L in sucrose gradient was compared between the control and *Ift20* mutant mice. After centrifugation, the collected gradients were analyzed by SDS-PAGE gels, and the presence of the two proteins was examined by specific antibodies. Both ODF2 and SPAG16L proteins were present in most gradients, from lighter density to greater density in the control. However, the two proteins were present only in the gradients with lighter density in mutant testes (Fig. 5F).

### Expression of autophagy core protein ATG16 is significantly reduced in the conditional *Ift20* mutant testis

IFT20 has been reported to play a role in autophagy, and it recruits autophagy core protein Atg16 to the autophagy core complex (Pampliega et al., 2013). Although some germ cells in the mutant testes completed all phases of spermatogenesis, the mutant spermatids were unable to process the redundant cytoplasm components of the cytoplasmic lobes for phagocytosis by the Sertoli cells, a sign of autophagy failure. Expression of Atg16 was significantly reduced in the conditional *Ift20* mutant mice compared to control (Fig. 5G).

## Discussion

Our laboratory identified a MEIG1/PACRG complex in male germ cells during the study of cargo transport along the manchette and sperm flagellum formation (Zhang et al., 2009; Teves et al., 2013; Li et al., 2015; Li et al., 2016). It appears that the roles of the MEIG1/PACRG complex and IFT proteins overlap in the transport of cargo during the assembly of cilia and flagella. Therefore, we decided to extend the study of MEIG1 to IFT and investigate the role of IFT in spermatogenesis and mammalian sperm flagella formation.

Even though mammalian male gametes have the longest flagella, very few studies have been conducted to investigate the role of IFT in the regulation of mammalian sperm flagellum formation. This is largely due to the fact that IFT is essential for embryonic development and complete knockout of an IFT component usually causes embryonic lethality (Nonaka et al., 1998; Marszalek et al., 1999).

The IFT20 expression pattern in male germ cells strongly suggests its role in cargo transport for building sperm flagella. Therefore, the floxed *Ift20* mice were crossed to the *Stra8-iCre* mice for the conditional knockout of *Ift20*. In *Stra8-iCre* transgenic mice, Cre recombinase (Cre) expression is directed by the *Stra8* (stimulated by retinoic acid gene 8) promoter fragment (Sadate-Ngatchou et al., 2008). This line has been shown to be useful in generating conditional knockouts in postnatal and premeiotic male germ cells for studying spermiogenesis (Bao et al., 2013).

The present study demonstrated that IFT20 is essential for normal spermatogenesis and male fertility. In the conditional *Ift20* mutant mice, very few germ cells completed spermatogenesis, which resulted in a significantly reduced sperm count. Histological examination of testis revealed normal mitosis and meiosis in the mutant mice. However, spermiogenesis, the final developmental phase was disturbed, which is consistent with the predicted role of IFT in carrying cargo for the formation of cilia and flagella. There are at least two explanations for the few spermatids that were able to fully differentiate and complete spermatogenesis. First, given that the deletion was not 100% in this model, the residual IFT20 protein might be sequestered in a few germ cells, through cytoplasmic bridges, at a level sufficient for select spermatid differentiation. Second, other IFT proteins may partially compensate for the loss of IFT20. Even though some mutant germ cells completed spermatogenesis, most had abnormal morphology and did not function well, as most sperm were immotile. Sperm motility in the mutant mice was significantly reduced compared to the control mice, and the sperm showed random movements instead of directional forward movement. The low sperm count and reduced sperm motility fully explain the infertility of adult conditional *Ift20* mutant males.

Why were a few of the younger mutant males subfertile at 6 weeks of age, but infertile after the first wave of spermatogenesis? It is likely the six mutant mice retained more IFT20 protein than the others that were completely infertile, but the protein level was still not enough for all germ cells to finish normal spermatogenesis, so fertility was reduced. After the first wave, less IFT20 protein was available, and the mutant mice completely lost fertility. Alternatively, other IFT proteins that could partially compensate for the loss of IFT20 may show differential expression during the first and subsequent waves of spermatogenesis. Also, it seems that while the first wave likely had relatively few normal or functional cells produced, these cells likely were passed along to the epididymis, but in subsequent waves there would be more cumulative or secondary effects on the germ cells and Sertoli cells would likely be responding to the abnormal germ cells and that would further reduce the chance of producing functional sperm.

The mutant abnormal sperm function seems to have been caused by disrupted sperm structure. Examination of epididymal sperm by light microscopy and SEM revealed multiple gross abnormalities, including short and kinked tails and round heads.

Interestingly, some sperm seemed to have redundant cytoplasm attached on the flagella, suggesting a failure to remove the cytoplasmic component of the residual bodies in mutant mice testes. TEM of testis and epididymal sperm further demonstrated abnormal spermiogenesis and sperm structure. Very few axonemes were discovered in seminiferous tubules of the mutant mice. Increased amounts of vesicles and fiber-like structures were discovered in the mutant cytoplasm, suggesting an impaired transport system in these cells and failure of incorporation of flagellar components into sperm tails. The significant number of sloughed germ cells in the lumen of seminiferous tubules in mutant mice also suggested a defect in Sertoli/germ cell recognition and failure of residual body formation and Sertoli cell phagocytosis of the excess germ cell cytoplasm.

Soon after the beginning of spermiation, a specialized structures, Tubulobulbar Complexes (TBCs) appear between the spermatid and the Sertoli cells (Vogl et al., 2014). These structures appear to be a modified form of endocytic machinery. Numerous proteins have been localized to specific sites within TBCs, including the integral membrane adhesion proteins nectin-2 (Sertoli cell) and nectin-3 (spermatid) (Guttman et al., 2004; Young et al., 2009), and associated vesicles that are positive for junction proteins also are positive for endosomal and lysosomal markers (Guttman et al., 2004; Young et al., 2009; Young et al., 2012). TBCs have been proposed to play a role in final sperm head and acrosome remodeling that occurs during spermiation, TBCs are thought to be involved in the reduction in the volume of the spermatid cytoplasm during spermiation (Russell, 1979; Russell, 1993). It remains to be determined if IFT20 is also involved in TBCs formation and function. Thus, it is reasonable to hypothesis that IFT20 would be involved in the transport of key proteins associated with TBC function.

In the few epididymal sperm of the mutant mice, many exhibited abnormal “9+2” axonemal structures, indicating improper flagella assembly, which is consistent with defective function of an IFT protein. Redundant cytoplasmic components were better illustrated by TEM when cross-sectional images were taken. It is believed that the retained cytoplasmic components mechanically obstructed the straightening of sperm flagella and affected their motility. This phenotype of failure to remove the residual body of leftover cytoplasm from late spermatids was previously reported in the *Sperm1* mutant mice (Zheng et al., 2007) and several other mutant mice, including *Tnp1*, *Tnp2*, *Prm1*, *Prm2*, *H1t2*, *Camk4*, or *Csnk2a2* (Xu et al., 1999; Wu et al., 2000; Cho et al., 2001; Zhao et al., 2001; Cho et al., 2003; Meistrich et al., 2003; Shirley et al., 2004; Zhao et al., 2004; Tanaka et al., 2005). It is not clear if IFT20 and these proteins function along similar pathways.

IFT proteins usually function as adaptors, as they associate with other proteins to form complexes for cargo transport (Lechtreck, 2015). Several IFT20 binding partners were identified, including IFT57, KIF3B, and CCDC41 (Sironen et al., 2010; Joo et al., 2013), but not ODF2 and SPAG16L, two flagellar proteins examined in this study. Given that localization of these two proteins in the elongating spermatids and epididymal sperm was dependent on IFT20, and their distribution in the sucrose gradient assay was shifted to the gradients with lower density, they appeared to be IFT20 cargo. However, IFT20 may not directly bind the two proteins, as other proteins may mediate the association. It is unlikely that the interaction was mediated through the known IFT20 binding partners; therefore, additional IFT20 binding partners might be present and should be identified in the future.

Even though IFT has been shown to be important for the assembly of cilia in most somatic tissues, it is dispensable for sperm flagellar assembly in both *Drosophila* and *Plasmodium falciparum*. It is thought that axonemal assembly occurs in the cytoplasm and the axoneme does not become membrane enclosed until after assembly (Han et al., 2003; Sarpal et al., 2003; Avidor-Reiss et al., 2004). However, the studies of *Ift88* and our conditional *Ift20* mutant mice revealed that IFT is essential for mouse spermiogenesis (San Agustin et al., 2015). Interestingly, ODF2 was affected in both *Ift88* and *Ift20* mutant mice, suggesting that different IFT proteins may transport the same cargo, or the IFT complex is disrupted by both mutations and the transport defect is secondary. We previously discovered that SPAG16L localization is also dependent on MEIG1/PACRG complex (Li et al., 2015). It is likely that the IFT20 and MEIG1/PACRG complexes function in the same transport pathway, although another possibility is that transporting one cargo may need multiple transport systems.

During spermiogenesis, the excess cytoplasmic organelles will need to be removed from the cell body. IFT20 not only regulates ciliogenesis through an IFT mechanism, it also associates with autophagy core protein Atg16L and helps to regulate a normal autophagy process required to clear redundant cytoplasmic components (Kobayashi, 2015). It is likely that IFT20 participates in autophagy found in the male germ cell, which involves the initiation and/or completion of the final steps required for the removal of late spermatid cytoplasmic residual body that is subsequently phagocytized by the Sertoli cells (Sprando and Russell, 1987c; Sprando and Russell, 1987a; Sprando and Russell, 1987b). This is supported by the following facts: a) the expression of the autophagy core protein Atg16L was significantly reduced in testes of the conditional *Ift20* mutant mice; b) sloughed residual bodies were present in the epididymal lumen; c) epididymal mutant sperm contained excess cytoplasmic components that were not removed during late spermiogenesis; and d) dramatically reduced mature lysosomes in the cytoplasm of the mutant mice. It has been shown that several autophagy core genes, such as *Atg7*, participate in the control of spermatogenesis (Wang et al., 2014). In medaka fish, *Ol-epg5* deficiency correlates with selectively impaired spermatogenesis and low allele transmission rates of the mutant allele caused by failure of germplasm and mitochondrial clearance during the process of germ cell specification and in the adult gonads (Herpin et al., 2015). Besides Atg16, IFT20 may also function through other autophagy core proteins. Thus, we assume that IFT20 forms at least two complexes; one for ciliogenesis and the other for autophagy. Since ciliogenesis and autophagy share similar molecular machinery (Kim et al., 2012; Diao et al., 2015; Liu et al., 2015), it is likely that the two IFT20 complexes have similar core proteins, while other proteins in the complexes determine the individual functions. For example, if the IFT20 complex associated with SPEF2, the complex would likely carry cargo for flagellogenesis; however, if the IFT20 complex associated with autophagy proteins, the complex would transport cargo for autophagy. Even though *Ift20* and *Ift88* mutant mice shared many common phenotypes, including reduced sperm count, disrupted flagellar structures, including disrupted axoneme and accessory structures, and ectopically assembled outer dense fibers and microtubules, suggesting that the two IFT proteins might function through a similar pathway in assembling sperm flagella. However, IFT20 mutant mice retained cytoplasmic lobe components that were not observed in IFT88 mutant mice suggesting that the two proteins have unique functions.

In summary, we studied the role of IFT20 in mouse sperm development and found evidence to support the hypothesis that IFT20 has at least two major functions: 1) transporting cargo such as ODF2 and SPAG16L to assemble the sperm flagellum, and 2) removing cytoplasmic components by initiating or helping to complete autophagy associated with the residual body. The molecular pathways involved with these two functions will need to be elucidated by identifying novel binding partners of IFT20.

## Materials and Methods

### Generation of Genetically Altered Mice

All animal work was approved by Virginia Commonwealth University's Institutional Animal Care & Use Committee (protocol AD10000167) in accordance with Federal and local regulations regarding the use of non-primate vertebrates in scientific research.

*Stra8-iCre* mice were purchased from Jackson laboratory (Stock No: 008208), and *Ift20*^*flox/flox*^ mice were generated previously (Jonassen et al., 2008). To obtain conditional *Ift20* mutant mice, *Stra8-iCre* males were bred with *Ift20*^*flox/flox*^ females. The resulting *Stra8-iCre*; *Ift20*^+/*flox*^ males were then crossed back with *Ift20*^*flox/flox*^ females. The conditional knockout mice (*Stra8-iCre*; *Ift20*^*flox/Δ*^) were considered as homozygous mutant mice, and *Stra8-iCre*; *Ift20*^+/*flox*^ mice were used as controls. To genotype mice, genomic DNA was extracted as described previously (Zhang et al., 2006). PCR was conducted using multiplex PCR mixture (Bioline, Cat No BIO25043). To examine *Stra8-iCre*, the following primers were used: Stra8-iCre forward: 5’-GTG CAA GCT GAA CAA CAG GA-3’; Stra8-iCre reverse: 5’-AGG GAC ACA GCA TTG GAG TC-3’. *Ift20* genotypes were identified as described previously (Jonassen et al., 2008).

## Assessment of Fertility and Fecundity

To test fertility, 6 weeks or older *Stra8-iCre*; *Ift20*^*flox*/ *Δ*^ (KO) and control males were paired with mature WT females for at least two months. Mating cages typically consisted of one male and one female. Mating behavior was observed and the females were checked for the presence of vaginal plugs and pregnancy. Once pregnancy was detected, the females were put into separate cages. The number of mice achieving a pregnancy and the number of offspring from each mating set or pregnancy were recorded.

## Western Blot Analysis

Testes were homogenized by ice-cold RIPA buffer (50 mM Tris-HCl [pH8.0], 170 mM NaCl, 5 mM ethylenediaminetetra-acetic acid, 1 mM dithiothreitol, and protease inhibitors [Complete mini; Roche Diagnostics GmbH]) containing either 1% NP-40, 0.5% sodium deoxycholate, and 0.05% SDS. After centrifuge 1200 rpm for 10 minutes, the supernatant was collected. Protein concentration was determined by Lowry assay. Denatured proteins were separated by SDS-PAGE and transfered to polyvinylidene fluoride membranes. Nonspecific sites were blocked with 5% nonfat dry milk in 0.5% Tween-20 in TBS for 1 h at room temperature, and the membranes was incubated overnight with indicated antibodies. After washing three times with TBS, the membranes were incubated with horseradish peroxidase-conjugated secondary antibodies (Amersham) for 2 hours at room temperature. After washing another three times by TBS, the membranes were visualized by less sensitive Pico or the higher sensitivity Femto detection system (Amersham Pharmacia) according to the manufacturer’s instructions.

## Transmission Electron Microscopy (TEM)

Mouse testes and epididymal sperm were fixed in 3% glutaraldehyde/1% paraformaldehyde/0.1 M sodium cacodylate, pH 7.4 at 4 °C overnight and processed for electron microscopy as reported. Images were taken with a Hitachi HT7700 transmission electron microscope.

## Scanning Electron Microscopy (SEM)

For SEM analysis, mouse epididymal sperm were collected and fixed in the same fixative solution as TEM. The samples were processed by standard methods (Jonassen et al., 2008). Images were taken with a Zeiss EVO 50 XVP SEM at Virginia Commonwealth University’s Microscopy Facility.

## Spermatozoa Counting

Sperm cells were collected from cauda epididymides and fixed in 2% formaldehyde for ten minutes at room temperature. Sperm were counted using a hemocytometer chamber under a light microscope and sperm number was calculated by standard method.

## Spermatozoa Motility Assay

Sperm were collected after swimming out from the cauda epididymides into PBS at 35°C for 10 min. The solution had a final pH of 7.35. Sperm populations were analyzed as soon as possible after their release from the epididymides. The sperm motility was observed using an inverted microscope (Nikon) equipped with 10X objective. Movies were recorded at 30 frames per second with SANYO color CCD, Hi-resolution camera (VCC-3972) and Pinnacle Studio HD (Ver. 14.0) software. For each sperm sample, a total of 10 fields were analyzed. Individual spermatozoa were tracked using the NIH ImageJ (NIH, Bethesda, MD, USA) and the plugin MTrackJ. Sperm motility was calculated as curvilinear velocity (VCL), which is equivalent to the curvilinear distance (DCL) that is travelled by each individual spermatozoon in one second (VCL=DCL/t).

## Histology and Immunofluorescence Staining on Tissue Sections

Testes and epididymides of adult mice were fixed in 4% formaldehyde solution in PBS, paraffin-embedded, and sectioned into 5 μm slides. For histology, Hematoxylin and Eosin staining on these sections was carried out using standard procedures. For immunofluorescence staining, testicular sections were incubated with an anti-IFT20 antibody, followed by a cy3 labeled goat anti-rabbit antibody. After washing three times in 1× PBS, each slide was mounted with VectaMount with DAPI (Vector Labs. Burlin-game, CA). Images were captured by confocal laser-scanning microscopy.

## Isolation of Spermatogenic Cells and Immunofluorescence(IF) Analysis

Testes from adult mice were dissected into a petri dish with 5 mL DMEM containing 0.5 mg/mL collagenase IV and 1.0 μg/mL DNAse I (Sigma-Aldrich), and was incubated for 30 min at 32°C with gentle stirring. The released spermatogenic cells were pelleted by centrifugation (5 min at 1000 rpm, 4°C). After washing with PBS, the cells were fixed in 5ml of 4% paraformaldehyde (PFA) containing 0.1M sucrose at room temperature for 15 minutes with constantly shaking. Dispersed mixed testicular cells were washed three times again with PBS. At the final wash, the cells were resuspended in 2 ml of PBS and 50 μl of cell suspension was loaded to slide and allowed to air-dry. To conduct IF, the cells were permeabilized with 0.1% Triton X-100 (Sigma-Aldrich) for 5 min at 37°C, washed with PBS three times, and blocked at room temperature for 1 hr with 10 % goat serum in PBS. Then the cells were washed three times in PBS and incubated overnight with indicated antibodies. After washing, the cells were incubated with the indicated secondary antibodies at room temperature for 1 h. The slides were then washed by PBS, mounted with VectaMount with DAPI (Vector Labs. Burlingame, CA), and sealed with nail polish. Images were captured by confocal laser-scanning microscopy as before.

## Sucrose gradient analysis of cytoplasmic extracts of testes

Adult control and *Ift20* mutant mice were killed by CO2 asphyxiation, and the testes were removed, decapsulated, dissected and homogenized as rapidly as possible. Cytoplasmic extracts were prepared by lysis of two testes in 0.5 ml HNM buffer (0.1 M NaCl, 3 mM MgCl_2_, 20 mM HEPES, pH 7.4) containing 0.5% Triton N-101 with 10 strokes of a motor driven glass–teflon homogenizer, and centrifugation at 13000 × g for 2 min. The supernatants were layered on a 5 ml 15–40% (w/w) linear sucrose gradient in HNM buffer, centrifuged at 28000 rpm for 150–175 min in the Beckman SW40 rotor, decelerated with the brake to 8000 rpm. The gradients were collected as seventeen 0.3 ml fractions (Cataldo et al., 1999).

## ACKNOWLEDGMENTS

We thank Dr Scott C. Henderson, Frances K. White, and Judy C. Williamson for their assistance with using the confocal microscopy and electronic microscopy in Microscopy Core Facility of Virginia Commonwealth University.

## Competing interests

The authors have no conflict of interest.

## Author contribution

Zhibing Zhang designed the experiments and wrote the paper. Zhengang Zhang., W. L., Y. Z., L. Z., M. E. T., H. L., J. L, and Zhibing Zhang performed the experiments. J. F S. edited the manuscript., G. J. P. contributed reagents and materials. J. A. F., R. A. H., and Zhibing Zhang analyzed the data.

## Funding

This research was supported by NIH grant HD076257, Virginia Commonwealth University Presidential Research Incentive Program (PRIP) and Massey Cancer Award (to ZZ), NIH grant GM060992 (to GJP), Chenery and Rashkind Grants from Randolph-Macon College (JF). Youth Talents of Science and Technology Projects of Health Department of Hubei Province of China (WJ2015Q026), Department of Hubei Province of China (QJX2012-22), Natural Science Foundation of China (81571428, 81502792, 81300536, 81172462).

Confocal microscopy and SEM were performed in the VCU Microscopy Facility of Virginia Commonwealth University (5P30NS047463). TEM was performed at Randolph-Macon College (NSF1229184).

